# *S*-Acylation of plant immune receptors mediates immune signaling in plasma membrane nanodomains

**DOI:** 10.1101/720482

**Authors:** Dongqin Chen, Nagib Ahsan, Jay J. Thelen, Gary Stacey

## Abstract

*S*-Acylation is a reversible protein post-translational modification mediated by Protein *S*-acyltransferases (PATs). Here, we demonstrate that three plant immune receptors P2K1 (DORN1), FLS2 and CERK1, representing three distinct families of receptor like-kinases, are *S*-acylated by *Arabidopsis* PAT5 and PAT9 on the plasma membrane (PM). Mutations in *Atpat5* and *Atpat9* resulted in an elevated plant immune response, increased phosphorylation and decreased degradation of FLS2, P2K1 and CERK1 during immune signaling. Mutation of key cysteine residues *S*-acylated in these receptors resulted in a similar phenotype as exhibited by *Atpat5*/*Atpat9* mutant plants. The data indicate that *S*-acylation also controls localization of the receptors in distinct PM nanodomains and, thereby, controls the formation of specific protein-protein interactions. Our study reveals that *S*-acylation plays a critical role in mediating spatiotemporal dynamics of plant receptors within PM nanodomains, suggesting a key role for this modification in regulating the ability of plants in respond to external stimuli.

## INTRODUCTION

Unlike mobile animals that contain specialized immune cells or organs that can trigger immune responses autonomously, plants are sessile organisms that require innate immunity to defend against detrimental microbes or pests. In order to perceive and trigger innate immune responses, plants employ a two-tier innate immune system, which includes critical plasma membrane (PM) localized receptors (Dodds and Rathjen, 2010). The first layer of defense is termed pattern-triggered immunity (PTI), since it relies on PM, pattern-recognition receptors (PRRs), which recognize conserved, pathogen-associated molecular patterns (PAMPs). Pathogens can also cause cell damage releasing damage-associated molecular patterns (DAMPs) that are also recognized by PM-localized, PRRs (Zipfel, 2014). PTI can be defeated by pathogen-produced, effector proteins that target a variety of cellular components involved in innate immunity. Hence, the second layer of defense, termed effector trigger immunity (ETI), involves plant recognition systems that counter the action of these pathogen effector proteins (Zhang et al., 2018).

PRRs encompass a variety of receptor-like kinase (RLK) subfamilies. Receptor-like kinases comprise a ligand-binding ectodomain, a transmembrane domain and an intracellular kinase domain. For example, FLS2, containing a leucine rich repeat (LRR) ectodomain, recognizes bacterial flagellin via direct binding to a conserved 22-aminoacid epitope, flg22 (Chinchilla et al., 2006; Gomez-Gomez and Boller, 2000; Zipfel et al., 2004). Another well-studied plant LRR-RLK is the *Arabidopsis* EFR receptor, which recognizes EF-Tu via perception of the conserved N-acetylated epitope elf18 (Kunze et al., 2004; Zipfel et al., 2006). A second family of RLKs involved in PAMP recognition is the lysin motif (LysM-RLK) family, which includes CERK1. Chitin is a major constituent of fungal cell walls, which is directly bound by AtCERK1 (AtLYK1) and AtLYK5 in *Arabidopsis* (Cao et al., 2014a; Miya et al., 2007). The first plant, DAMP PRRs identified (PEPR1 and PEPR2), members of the LRR-RLK subfamily, were shown to specifically recognize AtPEP peptides released due to cell/tissue damage (Yamaguchi et al., 2010; Yamaguchi et al., 2006). Extracellular ATP (eATP) is also released during tissue damage or in response to specific elicitation, including pathogens (Cao et al., 2014b). A member of the lectin-RLK subfamily, P2K1 (DORN1), was shown to be the key receptor that recognizes eATP resulting in the induction of an innate immunity response (Choi et al., 2014).

While different PRRs recognize different PAMPs/DAMPs, the plant responses induced upon activation of these receptors are highly similar. For example, PTI is characterized by an elevation of cytoplasmic calcium (Ca^2+^), reactive oxygen species (ROS), nitric oxide (NO), the activation of mitogen-activated protein kinases (MAPKs), and the expression of immune related genes (Cao et al., 2014b; Macho and Zipfel, 2014). Activation of the PRR by its cognate ligand leads to autophosphorylation and transphosphorylation of other proteins that is often followed by receptor ubiquitination and degradation, which subsequently dampens the immune response (Mithoe and Menke, 2018).

In contrast to phosphorylation, other forms of protein covalent modification are poorly studied in plants, although they are known to occur. For example, *S*-acylation, a reversible acylation of cysteine residues via a thioester linkage, is well characterized in mammals but poorly studied in plants. Mammalian studies have shown a critical role for *S*-acylation in controlling PM association, subcellular trafficking, stability, protein-protein interactions, enzymatic activity, and many other functions (De and Sadhukhan, 2018; Ko and Dixon, 2018; Sobocinska et al., 2018). Dynamic protein *S*-acylation is catalyzed by protein S-acyl transferases (PATs) that contain a conserved Asp-His-His-Cys (DHHC) catalytic domain, while deacylation is mediated by acyl protein thioesterases (APTs) (De and Sadhukhan, 2018). In human, a family of 24 DHHC-PATs was identified and shown to be involved in many physiological processes, as well as diseases spanning from neuropsychiatric disorders to cancers, cellular differentiation, melanomagenesis and so on (Chen et al., 2017b; De and Sadhukhan, 2018; Zhang and Hang, 2017). Most human DHHC-PATs are localized to endomembrane compartments such as Golgi, endosomes, endoplasmic reticulum, with only two proteins, DHHC20 and DHHC21, found at that the PM, which mediate epidermal growth factor receptor (EGFR) signaling in cancer and inflammatory responses (Beard et al., 2016; Runkle et al., 2016).

The genome of the genetic model plant, *Arabidopsis thaliana*, encodes 24 DHHC-PATs (Batistic, 2012). However, in contrast to humans, 12 *Arabidopsis* DHHC-PATs are localized to the PM, perhaps underlying the critical role that PM RLKs play in the ability of plants to respond to their environment (Batistic, 2012; De and Sadhukhan, 2018). There are relatively few studies describing the function of PATs in plants. Of the 24 encoded AtPATs, only six have been studied and shown to play a role in root hair growth (Hemsley et al., 2005; Wan et al., 2017), cell death, ROS production (Lai et al., 2015; Li et al., 2016), salt tolerance (Zhou et al., 2013), cell expansion and division (Qi et al., 2013), and branching (Xiang et al., 2010). Although a few plant proteins were previously shown to be *S*-acylated, including FLS2 (Hemsley et al., 2013), specific substrates for these AtPATs have not been identified, while their biological functions or mechanisms are also unknown.

In this manuscript, we demonstrate that two AtPATs, PAT5 and PAT9, can directly interact with and S-acylate three distinct plant PRRs, FLS2, CERK1 and P2K1, at the plasma membrane. There appears to be an inverse relationship between *S*-acylation and receptor activation, which is associated with autophosphorylation, in response to ligand addition. Concomitant with these changes, there is a marked relocalization of the receptor within the plasma membrane, which appears to govern protein-protein interactions within specific membrane nanodomains.

## RESULTS

### PAT5 and PAT9 restrict the plant immune response

In our previous study, we used a mass spectrometry-based *in vitro* phosphorylation strategy, termed kinase client assay (KiC assay) (Ahsan et al., 2013), to identify putative substrates of the P2K1 kinase domain; such as the NADPH oxidase respiratory burst oxidase homolog D (RBOHD) (Chen et al., 2017a). In addition, we found two homologous genes, *AtPAT5* and *AtPAT9*, which encode DHHC-PATs proteins likely involved in protein *S*-acylation (Figure 1A). Consistent with such a role, both PAT5 and PAT9 proteins were localized to the PM (Figure S1A).

**Figure 1.**
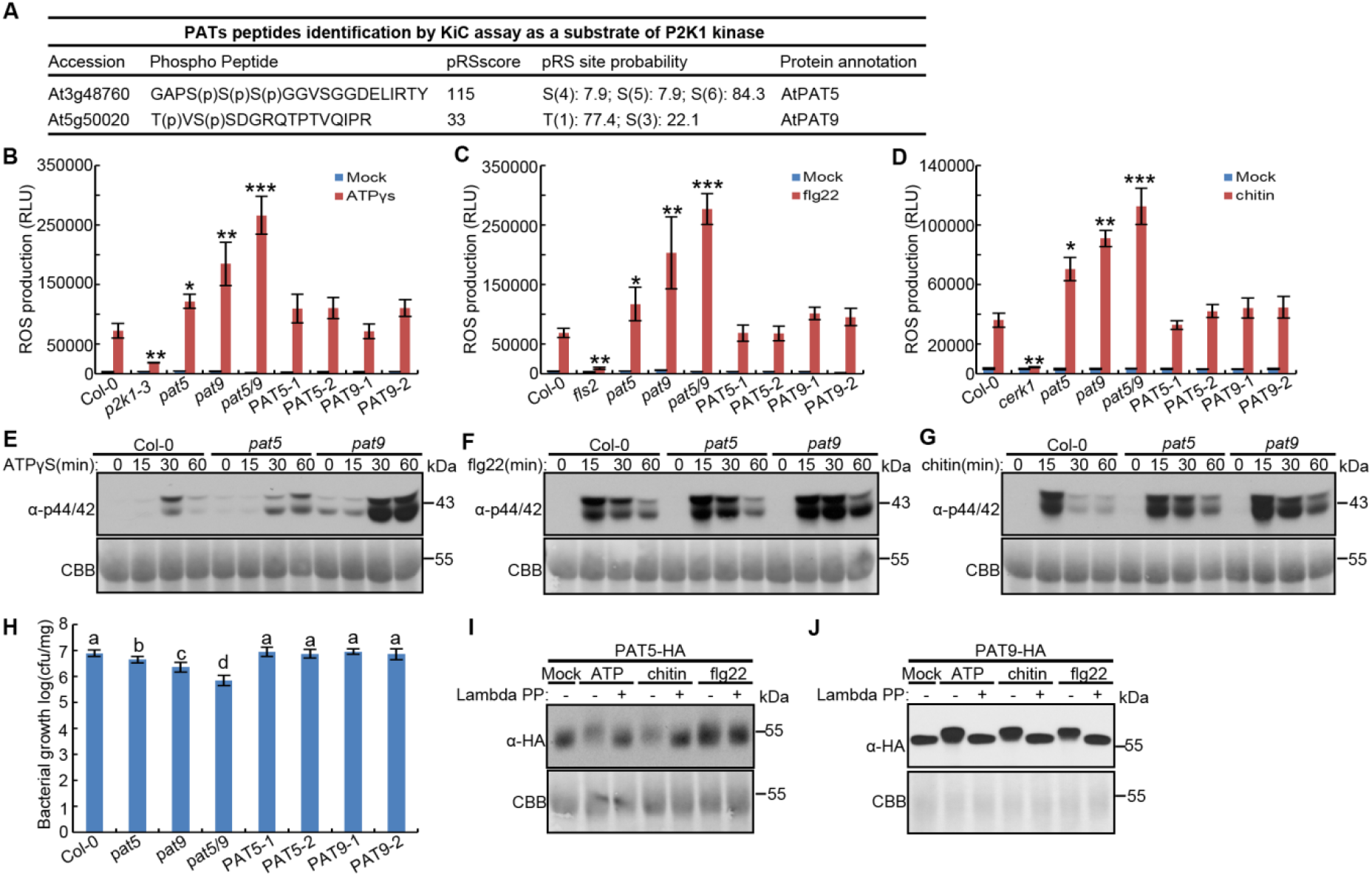
PAT5 and PAT9 control PTI response triggered by ATP, flg22 and chitin. (A) Identification of PAT5 and PAT9 tryptic peptides as a substrate of P2K1 kinase by KiC assay. (B-D) ROS production was measured in leaf discs after treatment with 200 μM ATPγS, 1 μM flg22 or 50 μg/ml chitin for 30 min. Leaf discs were taken from WT (Col-0), *pat5, pat9* and *pat5/9* double mutants, or their complemented lines *PAT5* (*NP::ATPAT5-HA/Atpat5*) and PAT9 (*NP::ATPAT5-HA/Atpat5*). RLU, relative luminescence units. (E-G) MAPKs activation of *Arabidopsis* leaf discs that treated with 200 μM ATPγS, 1 μM flg22 or 100 μg/ml chitin for the times indicated. Coomassie Brilliant Blue (CBB) staining of protein was used as loading control. (H) *Arabidopsis* seedlings with the indicated genotype (x axis) were inoculated with *Pst*. DC3000 bacteria and bacterial growth determined by plate counting 3 days post inoculation. (I and J) Ligand triggers PAT5 and PAT9 phosphorylation after treated with 100 μM ATP, 1 μM flg22 or 50 μg/ml chitin in their complemented lines *PAT5* (*NP::ATPAT5-HA/Atpat5*) and PAT9 (*NP::ATPAT5-HA/Atpat5*). Lambda protein phosphatase (Lambda PP, - and +) was added to release phosphate groups. CBB was used as loading control. For **B-D** and **H**, Error bars indicate ± SEM; n ⩾ 12 (biological replicates, one-sided ANOVA); means with different letters are significantly different; *P < 0.05, **P < 0.01, Student’s t test. See also figure S1.

Given their possible roles in modulating P2K1 receptor activities, we investigated the phenotype of plants defective in *Atpat5* or *Atpat9* by examining the response to different elicitors, including ATP, flg22 and chitin. Interestingly, the rapid elevation of cytoplasmic Ca^2+^ and ROS were significantly increased in both mutants compared to the wild type (Col-0) after ATP treatment (Figures 1B and S1B). Similar results were observed using flg22 or chitin as the elicitor (Figures 1C, 1D, S1C and S1D), suggesting that PAT5 and PAT9 might impact other PRRs. Consistent with the increase in cytoplasmic Ca^2+^ and ROS, ATP, flg22 or chitin-triggered MAPKs activation was also higher in the *Atpat5* and *Atpat9* mutants than Col-0, particularly at 30 and 60 min when wild type MAPK activation was decreasing (Figures 1E, 1F and 1G). Consistent with these stronger immune responses, growth of the bacterial pathogen *Pseudomonas syringae* DC3000 (*Pst*. DC3000) was significantly reduced in the *Atpat5* or *Atpat9* mutant plants (Figure 1H). These phenotypes were increased significantly in in *Atpat5/9* double mutant plants (Figures 1B, 1C, 1D, 1H, S1B, S1C and S1D), indicating that the functions of PAT5 and PAT9 are partially redundant.

In order to confirm PAT5 and PAT9 mediated PTI responses, we expressed the full-length PAT5 or PAT9 proteins, driven by their native promoters, in *Atpat5* and *Atpat9* mutants (Figure S1E), respectively. Expression of these proteins fully complemented the *Atpat5* and *Atpat9* mutant plants, restoring a wild-type PTI response (Figures 1B, 1C, 1D, 1H, S1B, S1C and S1D). Furthermore, we tested whether the activation of the PAT5 and PAT9 proteins was associated with those PTI responses. Indeed, addition of ATP, flg22 or chitin triggered phosphorylation of PAT5 and PAT9 (Figure 1I and 1J). Taken together, the above findings reveal that PAT5 and PAT9 are functional redundant and function to negatively regulate plant innate immunity mediated by at least three, distinct PRRs (FLS2, P2K1 and CERK1).

### PAT5 and PAT9 directly interact with and are phosphorylated by FLS2, CERK1 and P2K1

To verify the KiC assay result that PAT5 and PAT9 are kinase substrates of the P2K1 receptor (ATP-elicited PTI), we used a variety of assays to look for protein-protein interaction. These same assays were also used to test whether PAT5 and PAT9 were also kinase substrates of FLS2 (flg22-elicited PTI) or CERK1 (chitin-elicited PTI). For example, split-luciferase complementation imaging (LCI) assays in *Nicotiana benthaminana* showed that both PAT5 and PAT9 can interact *in vivo* with unstimulated P2K1, FLS2 and a kinase dead version CERK1 (D441A), but not RBOHC (Figures S2A). Interestingly, all these interactions were reduced 15 min after elicitation but enhanced 1 hour after elicitation (Figures S2A). Similarly, bimolecular fluorescence complementation (BiFC) assays in *Arabidopsis* protoplasts were used to specifically test for interactions at the PM. The yellow fluorescence signals were produced in the combination of PAT5/9-YFP^C^ with P2K1/FLS2/CERK1-YFP^N^ and merged well with the PM marker FM4-64 signal (Figure S2B). Interestingly, PAT5 and PAT9 appeared to form a heterodimer and homodimer in both assays (Figures S2A, S2B and S5C). These results indicate that PAT5 and PAT9 specifically associate with P2K1, FLS2, and CERK1 on the PM in a DAMP/PAMP-elicited manner.

The LCI and BiFC results show that PAT5/PAT9 and receptors can form a complex, but do not indicate that these proteins can interact directly and which domains are responsible for their interaction. Therefore, we performed *in vitro* pull-down assays using GST- or His-tagged recombinant proteins purified from *E. coli*. The PAT5 and PAT9 proteins are composed of four transmembrane domains (TMDs), the catalytic DHHC-Cysteine rich domain (DHHC-CRD) which is present in the cytosol and necessary for auto-acylation and for the modification of target proteins, the N- and C-terminal cytoplasmic domains (Figure S2C). Remarkably, PAT5-CRD, PAT5-C, PAT9-CRD and PAT9-C were able to directly bind to C-terminal cytoplasmic domains P2K1-KD, FLS2-CD and CERK1-C (Figure 2A). Meanwhile, an interaction with the kinase inactive form of the P2K1 kinase domain (P2K1-KD-1) was also observed in this pull-down assay (Figure 2A). These findings, together with the interaction results using the kinase dead version of CERK1^D441A^ in the LCI assays (Figure S2A), suggest that PAT5/PAT9-P2K1/CERK1 interactions are independent of receptor autophosphorylation. Moreover, the PAT5-P2K1 and PAT9-P2K1 interactions were also detected in Co-immunoprecipitation (Co-IP) assays when PAT5-Myc or PAT9-Myc were co-expressed with P2K1-HA in wild-type protoplasts (Figure 2B). The endogenous FLS2 and CERK1 proteins were detected in Co-IP experiments using transgenic plants carrying PAT5-HA and PAT9-HA under the control of their native promoters (Figures 2C and 2D). Consistent with the LCI results, the PAT5/PAT9-P2K1/FLS2/CERK1 interactions were minimal 5 min after elicitation but then markedly increased 30 min after elicitation with either ATP, flg22 or chitin (Figures 2B, 2C and 2D). Taken together, these various assays indicate that PAT5 and PAT9 directly interact with P2K1, FLS2 and CERK1 in a ligand-enhanced manner.

**Figure 2.**
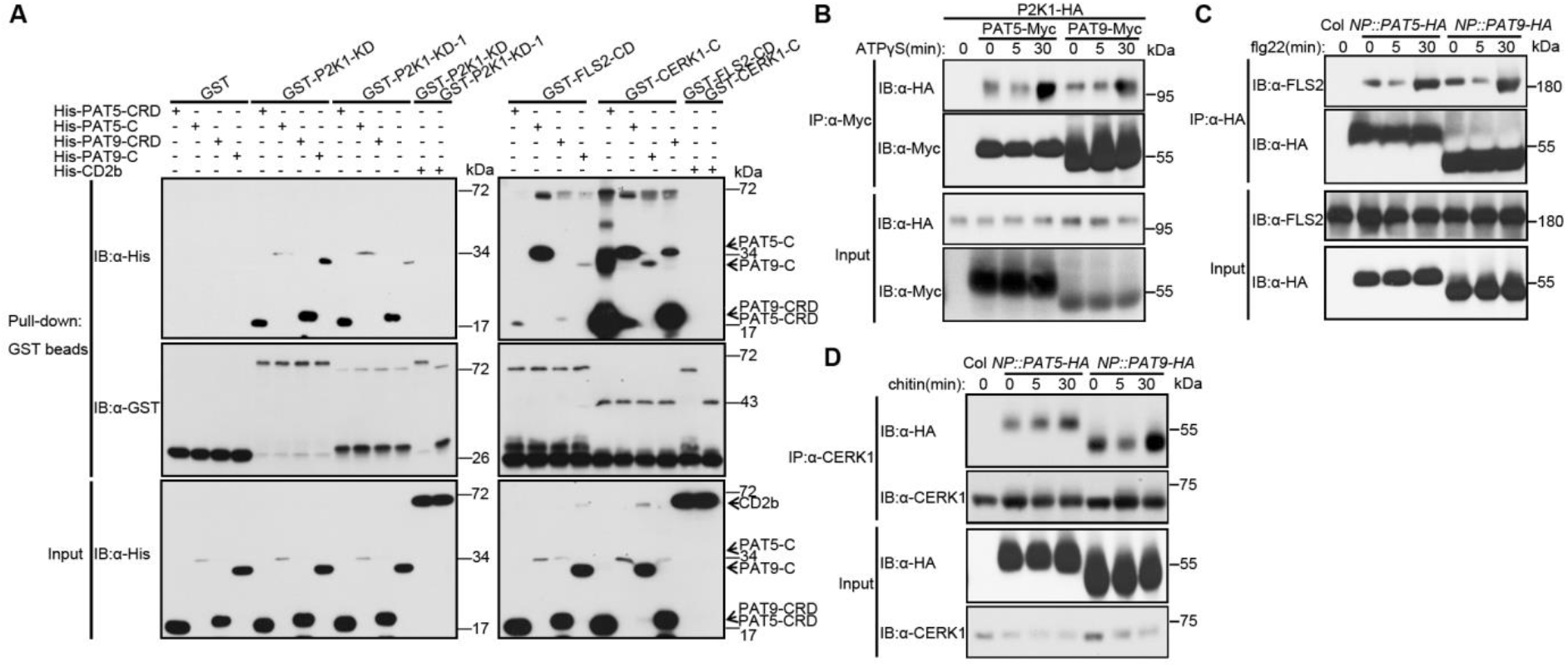
PAT5 and PAT9 directly interact with P2K1, FLS2 and CERK1 *in vitro* and *in vivo*. (A) Directly interaction of PATs and receptors *in vitro*. Purified, recombinant proteins were incubated with Glutathione Sepharose 4B beads followed by GST-His pull-down assay. His-CD2b (At5g09390, CD2-binding protein-related, a substrate of P2K1 from KiC assay data) was used as a negative control. (B) PAT5 and PAT9 interact with P2K1 *in vivo* in an ATP-enhanced manner. The indicated constructs were transiently expressed in *Arabidopsis* wild-type protoplasts treated with 50 μM ATPγS followed by Co-IP assays. (C and D) PAT5 and PAT9 interact with FLS2 and CERK1 in *Arabidopsis* plants. Stable *Arabidopsis* transgenic plants expressing *NP::ATPAT5/Atpat5* or *NP::ATPAT9/Atpat9* were treated with 1 μM flg22 or 50 μg/ml chitin and total protein extract was subjected to Co-IP. See also figure S2.

As the peptides of PAT5 and PAT9 were identified as the substrates of the P2K1 kinase (Figure 1A), we sought to determine whether FLS2 or CERK1 can also use these peptides as substrates. Consistent with the KiC assay results, P2K1-KD strongly trans-phosphorylated PAT5-CRD, PAT5-C, PAT9-CRD and PAT9-C, whereas the kinase dead P2K1-KD-1 failed to phosphorylate them (Figure S2D). Similar results were also observed using the CERK1-KD kinase domain, but not FLS2-KD, which showed low autophosphorylation ability (Figure S2E). The above data, together with PTI-induced phosphorylation of the PAT5 and PAT9 proteins, indicate that receptors can directly phosphorylate PAT5 and PAT9 in a DAMP/PAMP-induced manner.

### FLS2, CERK1 and P2K1 are *S*-acylated by PAT5 and PAT9

To investigate whether FLS2, CERK1 and P2K1 can be *S*-acylated and which residues are targeted, we first predicted the potential *S*-acylation residues using GPS-lipid 1.0 software (online http://lipid.biocuckoo.org/webserver.php) (Figure S3). The FLS2 predicted sites were the same as previously described (Hemsley et al., 2013). Based on these predictions, we generated a series of mutant forms of FLS2, CERK1 and P2K1 in which cysteine (C) sites were individually mutated to serine (S). These mutant proteins were then transiently expressed in *Arabidopsis* protoplasts and their *S*-acylation status determined through a parallel *S*-acylation assay with or without the hydroxylamine thioester-cleavage step. Compared with the wild-type FLS2, CERK1 and P2K1 forms, the levels of *S*-acylation from single mutants-such as P2K1^C394S^, P2K1^C407S^, CERK1^C381S^ and CERK1^C572S^ were significantly reduced, while their double mutants include FLS2^C830/831S^, p2K1^C394/407S^ and CERK1^C381/572S^ were below the level of detection (Figures 3A, 3B and 3C). Remarkably, P2K1^C394/407S^, CERK1^C381S^, CERK1^C572S^ and CERK1^C381/572S^ proteins migrated faster than the wild-type P2K1 and CERK1 when separated by SDS-PAGE (Figures 3B and 3C). However, the FLS2^C830/831S^ did not display such a protein mobility shift due to its high protein size (Figure 3A). Furthermore, loss of PAT5 and PAT9 function remarkably reduced FLS2, CERK1 and P2K1 *S*-acylation levels, while complemention by expression of the wild-type PAT5 or PAT9 proteins restored these deficiencies (Figures 3D, 3E and 3F). However, the PAT5^C188S^ and PAT9^C166S^ mutants, in which the DHHC catalytic domain was mutated to DHHS, lost auto *S*-acylation activity and failed to rescue FLS2, CERK1 and P2K1 *S*-acylation levels (Figures 3D, 3E and 3F) when expressed in the *Atpat* mutant background.

**Figure 3.**
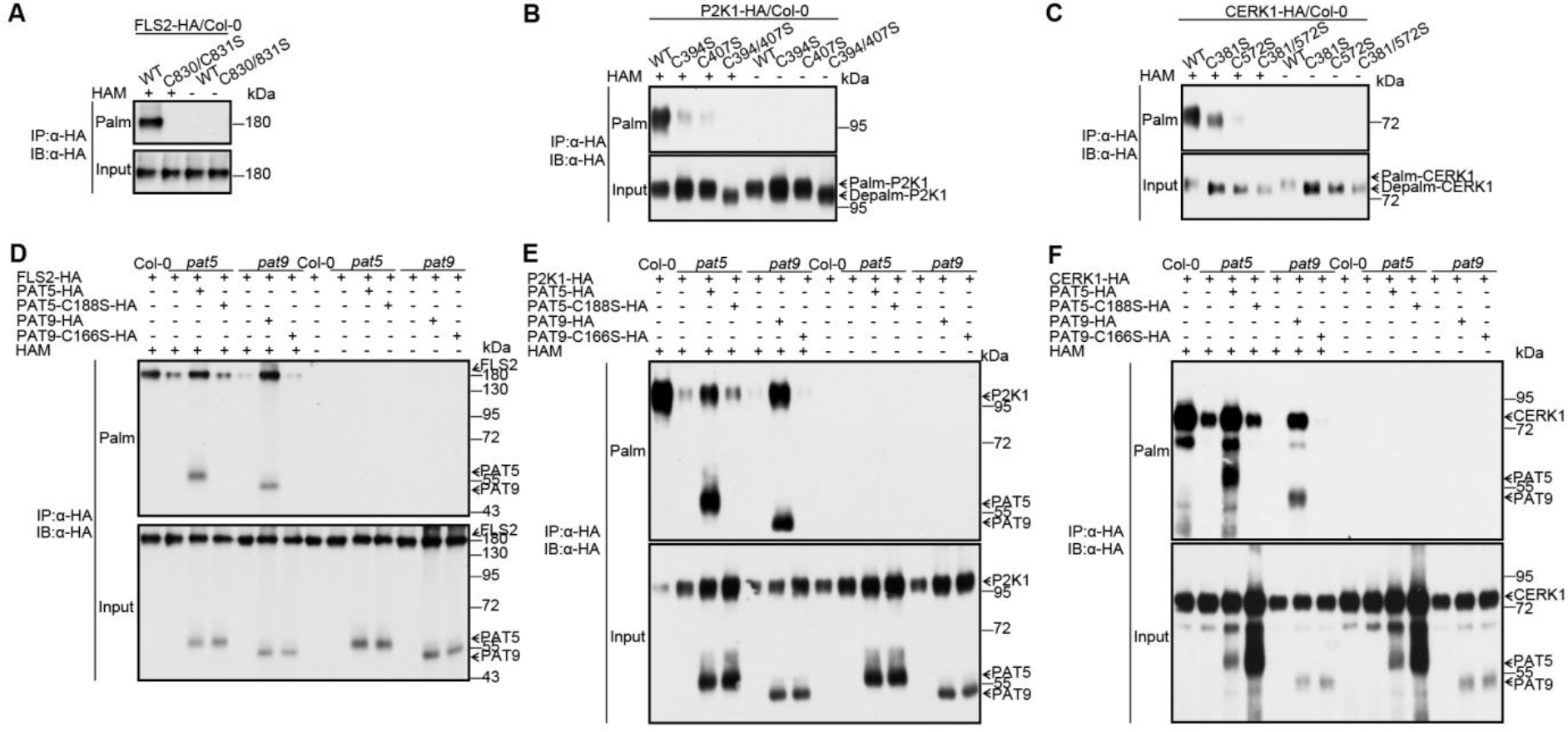
PAT5 and PAT9 *S*-acylate P2K1, FLS2 and CERK1. (A-C) Detection of P2K1, FLS2 and CERK1 *S*-acylation and their associated residues. The indicated constructs with HA tag were transiently expressed in *Arabidopsis* Col-0 protoplasts. The *S*-acylation levels were detected by an acyl-resin capture (acyl-RAC) assay. Palm-, palmitoylated protein; Depalm-, depalmitoylated protein. HAM, hydroxylamine, for cleavage of the Cys-palmitoyl thioester linkages. (D-F) PAT5 and PAT9 *S*-acylate P2K1, FLS2 and CERK1 *in planta*. Protoplasts of *Arabidopsis* wild-type Col-0, *Atpat5* and *Atpat9* mutants were transfected with plasmids as indicated, followed by acyl-RAC assays. See also figure S3.

The above data clearly indicate that PAT5 and PAT9 *S*-acylate FLS2, CERK1 and P2K1 *in planta*.

### *S*-Acylation mediates FLS2, CERK1 and P2K1 phosphorylation and degradation in a ligand-dependent manner

Given that PAT5 and PAT9 directly interact with, are phosphorylated by and *S*-acylate P2K1, FLS2 and CERK1 in a DAMP/PAMP-dependent manner, we sought to determine whether the activation of P2K1, FLS2 and CERK1 were regulated by PAT5 or PAT9. Previous results showed that autophosphorylation of P2K1 is induced by ATP elicitation (Chen et al., 2017a). Therefore, we expressed P2K1-HA in the *Atpat5* or *Atpat9* mutant background and found that, compared with *P2K1-HA*, the *P2K1-HA/Atpat5* and *P2K1-HA/Atpat9* plants showed greater P2K1 phosphorylation 15 and 30 min after ATP treatment (Figures 4A and 4B). The mobility of the phosphorylated protein was shifted by treatment with lambda protein phosphatase (lambda PP) to release any phosphate groups. On the other hand, the addition of 2-bromopalmitate (2-BP), a palmitate analogue known to inhibit *S*-acyl transferase activity, also increased P2K1 phosphorylation (Figure S4A). Ubiquitin-mediated degradation of FLS2 is ligand, dose, and time dependent (Smith et al., 2014). Indeed, endogenous FLS2 protein levels were significantly decreased 15 min after treatment with a high concentration of 20 μM flg22 in wild-type Col-0 plants. However, FLS2 degradation was significantly reduced in the *Atpat5* and *Atpat9* mutants or when wild-type plants were treated with 2-BP (Figures 4C and S4B). We also used α-CERK1 antibody to detect turnover of the endogenous CERK1 protein in Col-0 plants, which showed significant chitin-induced phosphorylation and degradation of CERK1 within 5 min of elicitation. In contrast, the level of endogenous CERK1 phosphorylation was significantly increased with a reduction in CERK1 degradation in the *Atpat5* or *Atpat9* mutant plants (Figure 4D). Moreover, these same effects were seen upon treatment with 2-BP (Figure S4C). These data clearly demonstrate that PAT5 and PAT9 modulate activation of P2K1, FLS2 and CERK1 protein in a ligand-induced manner, mediated by effects on both autophosphorylation and subsequent protein degradation.

**Figure 4.**
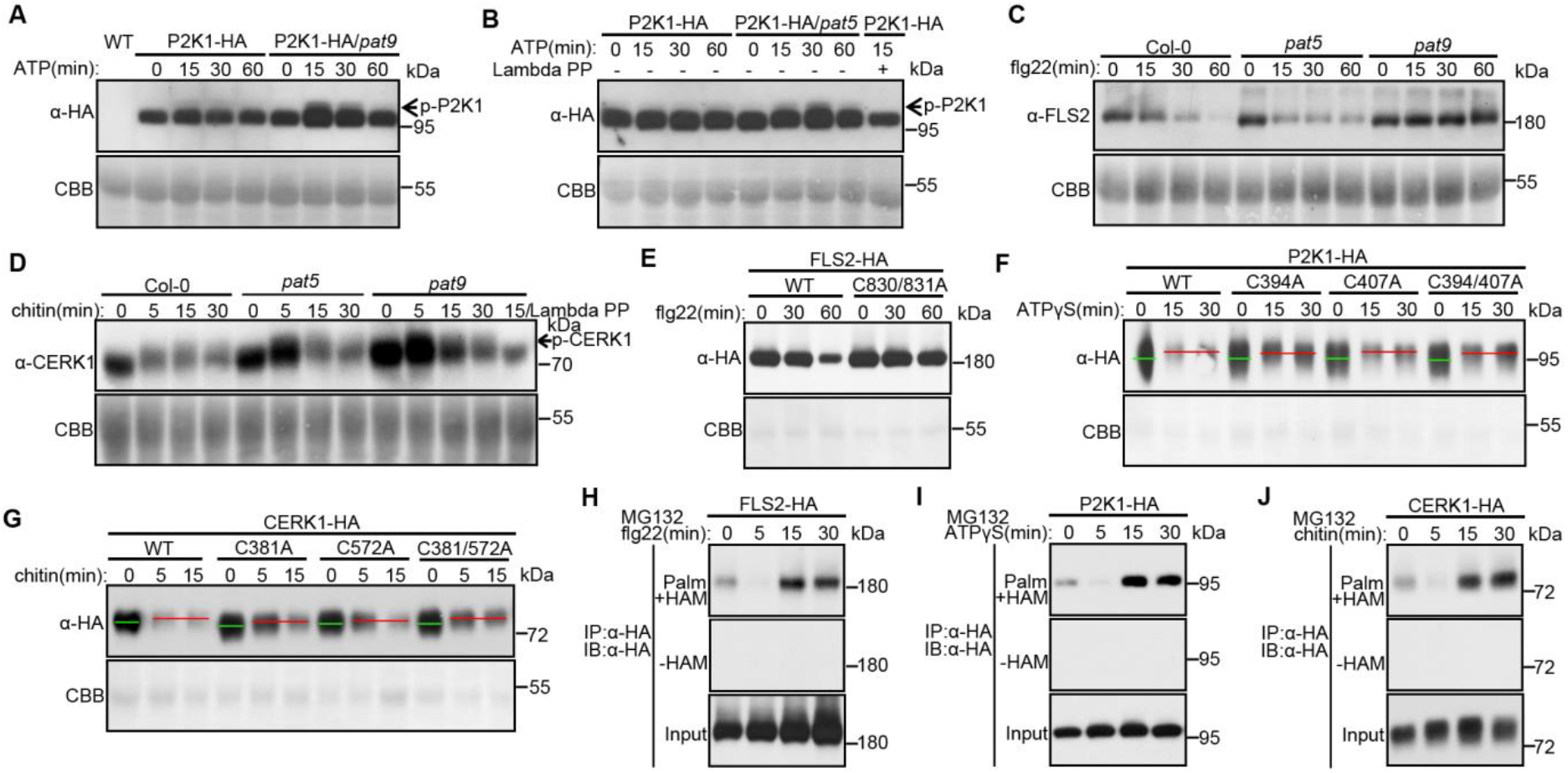
PAT5 and PAT9 regulate P2K1, FLS2 and CERK1 phosphorylation and degradation. (A and B) P2K1-HA phosphorylation was analyzed by immunoblot in wild type, *Atpat5* and *Atpat9* mutant backgrounds upon addition of 200 μM ATPγS. Lambda PP was added to release phosphate groups. p-P2K1, phosphorylation of P2K1. (C) flg22-induced endogenous FLS2 degradation in leaf discs of Col-0, *Atpat5* and *Atpat9* mutant plants after treatment with 20 μM flg22. (D) The endogenous CERK1 phosphorylation and degradation were detected in leaves of Col-0, *Atpat5* and *Atpat9* mutant plants infiltrated with 100 μg/ml chitin. p-CERK1, phosphorylation of CERK1. (E-G) Analysis of the rate of turnover of P2K1, FLS2 and CERK1 proteins modified (C→A) at the site of *S*-acylation. The indicated constructs were expressed into *Arabidopsis* wild-type Col-0 protoplasts treated with 250 μM ATPγS, 10 μM flg22 or 100 μg/ml chitin. Green lines, dephosphorylated proteins; Red lines, phosphorylated proteins. Proteins were separated by high-resolution SDS-PAGE. CBB was used as loading control. (H-J) Dynamic *S*-acylation levels of FLS2, P2K1 and CERK1 in stable transgenic seedlings upon 100 μM ATPγS, 1 μM flg22 or 50 μg/ml chitin. MG132 was used to inhibit receptors degradation. See also figure S4.

In order to better understand how *S*-acylation mediates FLS2, CERK1 and P2K1 activity upon elicitor stimulation, we transiently expressed mutated receptors proteins in *Arabidopsis* protoplasts. Expression of the FLS2^C830/831S^, P2K1^C394S^ and CERK1^C381/572S^ proteins showed less protein degradation than the wild-type forms upon flg22, ATP and chitin treatment, respectively (Figures S4D, S4E and S4F). The substitution of C residue with a S residue could provide a potential phosphorylation site and, therefore, we also replaced each C residue with alanine. The alanine substitute forms of each of the proteins (e.g., FLS2^C830/831A^) also significantly reduced ligand-induced protein degradation (Figure 4E). Meanwhile, elicitor-induced protein phosphorylation of the P2K1^C394A^, P2K1^C407A^, P2K1^C394/407A^, CERK1^C381A^, CERK1^C572A^ and CERK1^C381/572A^ proteins were significantly increased, while protein degradation were remarkably decreased (Figures 4F and 4G). In summary, these results further demonstrate that *S*-acylation of key residues of the FLS2, CERK1 and P2K1 proteins plays an important role in mediating ligand-induced autophosphorylation and degradation.

In order to better examine the dynamic effects of *S*-acylation in the absence of receptor degradation, we treated plants with MG132, an inhibitor of proteasome-mediated protein degradation. Under these conditions, *S*-acylation of FLS2-HA, P2K1-HA or CERK1-HA was significantly decreased 5 min after ligand addition but remarkably increased 15 and 30 min after elicitation (Figures 4H, 4I and 4J), showing a clear inverse relationship to the levels of autophosphorylation of the receptor proteins (Figures 4A and 4D). These results, together with the data of immune response (Figure 1), indicate that *S*-acylation is first abolished by ligand-induced receptor activation, but then restored concomitant with an abatement of immune signaling.

### *S*-acylation regulates FLS2, CERK1 and P2K1 localization into PM microdomains

As mentioned previously, studies in mammalian cells showed that *S*-acylation is a key factor in subcellular trafficking or membrane association of key receptor proteins, including specific localization into lipid rafts (PM nanodomains) (De and Sadhukhan, 2018). A similar phenomenon was suggested by the observation of dispersed punctate structures with increased fluorescence intensities when we conducted the BiFC assays and examined them using confocal laser scanning microscopy (Figures 5A and S5A). We therefore perfected the single-particle tracking assay using GFP- and RFP-tagged fluorescent proteins in *Arabidopsis* protoplasts. Specifically, FLS2-, CERK1- and P2K1-GFP proteins localized to PM nanodomains and co-localized with the protein marker FLOT1-RFP (Figures 5D and S5B). Flotillin (FLOT1) is a known marker for PM lipid rafts, and has, for example, been implicated in the endocytosis of FLS2 in response to flg22 (Yu et al., 2017). Meanwhile, LCI assays showed that PAT5, PAT9, FLS2, CERK1 and P2K1 could also interact with FLOT1 *in vivo* (Figure S5C), suggesting that FLOT1 may be involved in ligand-activated endocytosis of a wide range of PM receptor proteins.

**Figure 5.**
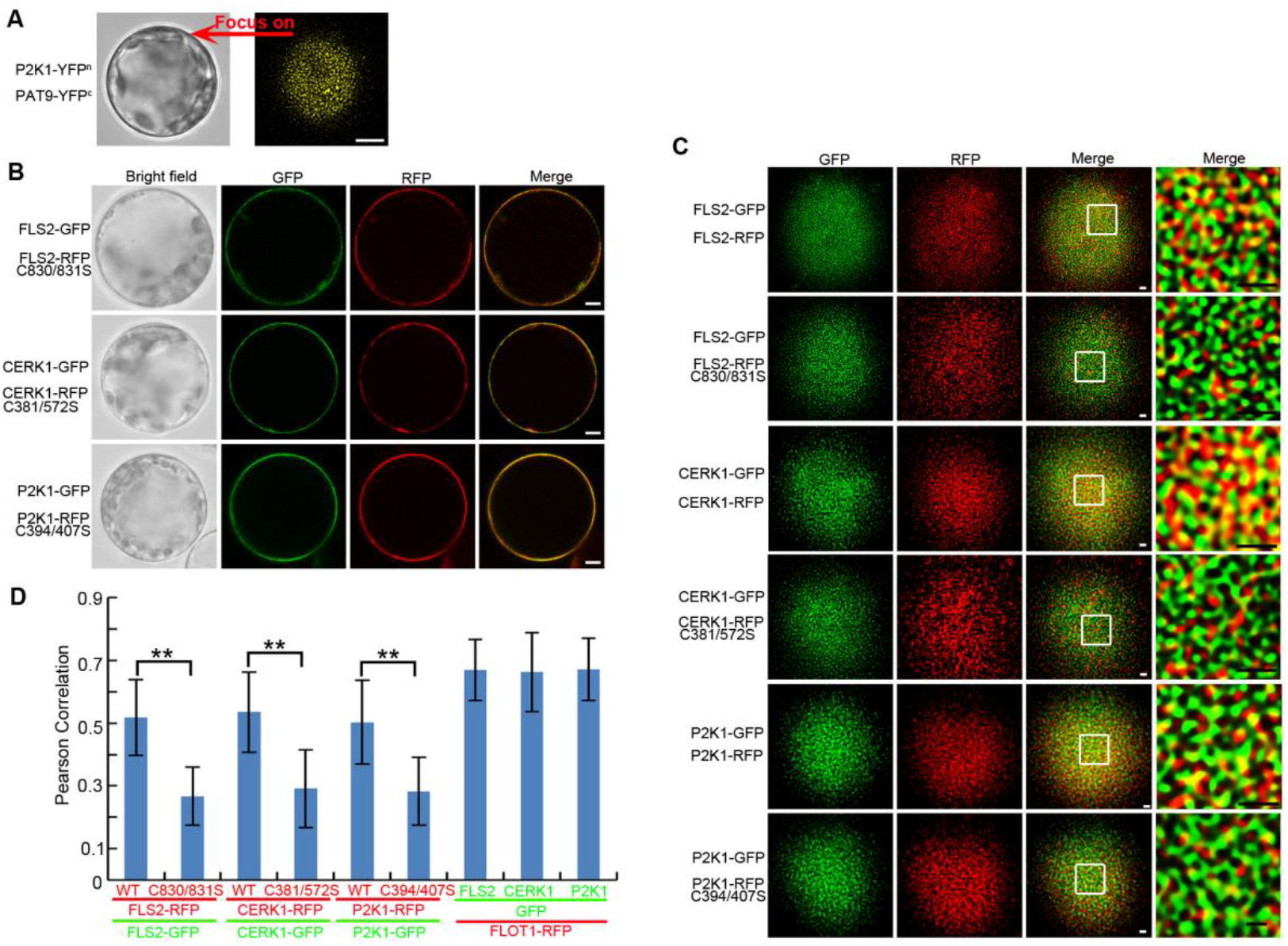
*S*-acylation mediates localization of FLS2, CERK1 and P2K1 into PM nanodomains. (A) Dispersed punctate fluorescence signaling was observed by focusing the fluorescent microscope on the top of *Arabidopsis* protoplasts in BiFC assay. Bar = 5 μM. (B) The *S*-acylation state of FLS2, CERK1 or P2K1 does not affect their trafficking to the PM. Bar = 5 μM. (C) *S*-Acylation regulates localization of FLS2, CERK1 and P2K1 to PM nanodomains. The indicated constructs were expressed in *Arabidopsis* wild-type Col-0 protoplasts, followed by confocal microscope imaging. The dashed squares represent the areas magnified within the far right image. Bar = 1 μM. (D) Quantitative co-localization analysis of the confocal micrographs shown in panel **C** using Pearson correlation. Error bars indicate ± SD; n = 8 (biological replicates, one-sided ANOVA); **P < 0.01, Student’s t test. See also figure S5.

Next, we tested the specific role that *S*-acylation might play in PM localization. We examined the three *S*-acylation mutants of FLS2^C830/831S^, CERK1^C381/572S^ and P2K1^C394/407S^, which showed identical PM localization to that of the FLS2, CERK1 and P2K1 proteins (Figure 5B). These data indicate that three receptors are correctly trafficked to the plasma membrane in the absence of *S*-acylation. Hence, unlike the case of G-protein-coupled or glutamate receptors in mammals (Zareba-Koziol et al., 2018), *S*-acylation is not required for correct PM targeting of these proteins. We next asked the question of whether the three PRRs localized to the same PM nanodomains. Indeed, examination of the different wild-type combinations of FLS2-GFP/FLS2-RFP, CERK1-GFP/CERK1-RFP and P2K1-GFP/P2K1-RFP showed that they co-localized into the same PM nanodomains (Figure 5D). In contrast, the FLS2-GFP/FLS2^C830/831S^-RFP, CERK1-GFP/CERK1^C381/572S^-RFP and P2K1-GFP/P2K1^C394/407S^-RFP combinations showed a marked reduction in co-localization compared with the wild-type receptor combinations (Figure 5C). In order to quantify their co-localization, we determined Pearson correlation coefficients of the images, which is a measure of the linear correlation between two variables X and Y. Our quantitative co-localization analysis revealed that co-localization of the mutant proteins, unable to be *S*-acylated, had significantly lower Pearson correlation coefficients (Figure 5D). Hence, the data indicate that the various PRRs co-localize to PM nanodomains with the known lipid raft protein, flotillin, and that this PM location is dependent on the level of PAT5/9 mediated *S*-acylation of the receptor proteins.

### FLS2, CERK1 and P2K1 *S*-acylation regulates PTI

In order to demonstrate the biological relevance of CERK1 and P2K1 *S*-acylation mediated by PAT5 and PAT9, we generated stable transgenic plants expressing the mutated, *S*-acylation site proteins in the respective *cerk1*, or *p2k1-3* mutant backgrounds under the control of their native promoters (Figure S5A and S5B). Similar to the data derived using the *Atpat5* and *Atpat9* mutant plants, ligand-induced elevation of cytoplasmic calcium levels in plants expressing P2K1^C394A^, P2K1^C407A^, P2K1^C394/407A^, CERK1^C381A^, CERK1^C572A^ or CERK1^C381/572A^ variants were significantly higher than similar mutant plants complemented with the corresponding wild-type gene (Figures S6C and S6D). Notably, the resistance to the virulent bacterium *Pst*. DC3000 *hrcC*^−^ was also significantly enhanced in plants expressing FLS2^C830/831A^, P2K1^C394A^, P2K1^C407A^, P2K1^C394/407A^, CERK1^C381A^, CERK1^C572A^ or CERK1^C381/572A^ compared to the corresponding wild-type controls (Figures 6A and 6B). In summary, consistent with both the *in vivo* and *in vitro* biochemical measurements, *S*-acylation of FLS2, CERK1 or P2K1 negatively regulates the elicitor mediated PTI responses.

**Figure 6.**
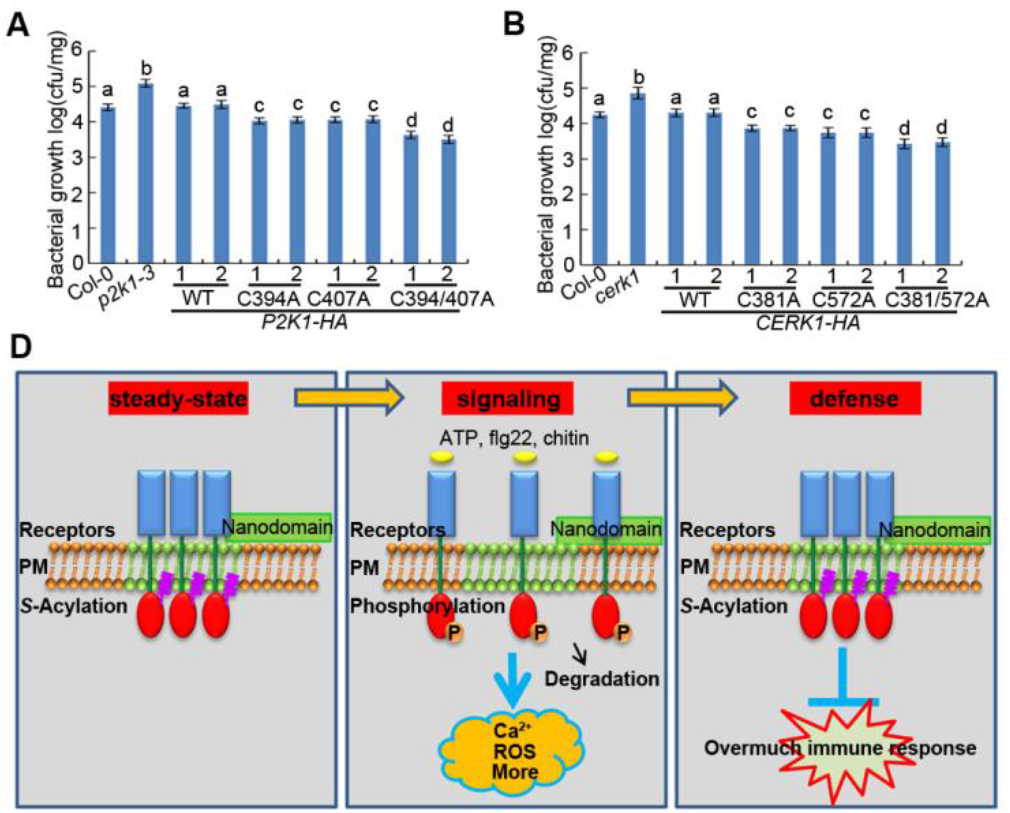
*S*-Acylation of FLS2, CERK1 and P2K1 negatively regulates PTI. (A and B) The indicated seedlings were flood-inoculated with *Pst*. DC3000 *hrcC*^−^ bacteria and bacterial growth measured by plate counting after 2 days post inoculation. Values represent the mean ± SEM, n ⩾ 12 (biological replicates); Means with different letters are significantly different (P < 0.05; one-sided ANOVA). (C) Model for role of receptor *S*-acylation in PM localization and signaling. In the absence of an inducing ligand (steady-state), plant receptors are *S*-acylated and inactivated, localized within discrete PM nanodomains. Upon addition of the activating ligand, the receptors are rapidly deacylation, altering the stability of the nanoclusters, with subsequent autophosphorylation, and downstream signaling leading to PTI. Activation of the receptors also triggers their ubiquitin-mediated degradation. After defense events, *S*-acylation inactivates receptors to PM nanodomains and dampens the immune response to protect plant growth. See also figure S6.

## DISCUSSION

Dynamic *S*-acylation determines protein function by influencing their association with membranes, compartmentalization in membrane domains, trafficking, and stability (Sobocinska et al., 2018). In humans, many *S*-acylated receptors are found in membrane nanodomains (lipid rafts), originally characterized as low density detergent-resistant membrane domains, by sucrose density gradient centrifugation isolation (Poggi et al., 2013; Sobocinska et al., 2018). However, the recent advent of high resolution microscopy has moved the identification of such structures beyond such an empirical definition allowing direct observation and study (Lu and Fairn, 2018).

Unlike animals, plants are unable to move in response to danger or stress nor do they possess humoral and cellular immunity. Instead, plants have evolved an extensive and complex system of receptors that recognize pathogens, pests and cellular damage through a large and diverse family of PM-localized RLKs. Ligand activation of these receptors leads to autophosphorylation, transphosphorylation of downstream proteins, and other cellular events culminating ultimately into a robust innate immunity response (Monaghan and Zipfel, 2012). Concomitant with these events, the receptor is rapidly turnover through endocytosis, presumably to dampen the immune response that, while protective, is also detrimental to overall plant growth. Although each PRR recognizes a specific PAMP/DAMP signal, in general, the downstream signaling responses are highly similar regardless of which PRR is activated. Some receptors, such as BRASSINOSTEROID INSENSITIVE 1 (BRI1) and FLS2 are similar but mediate quite distinct responses; that is, brassinosteroid growth responses (BRI1) and PTI (FLS2) (Bucherl et al., 2017). It was suggested that the specificity of these receptors is mediated by their specific localization in PM nanoclusters with BRI1 and FLS2 showing distinct and separate localization (Bucherl et al., 2017). However, little is known as to how such localization may be mediated or what is required to co-localize receptors and essential, accessory proteins into such nanodomains.

The current study demonstrates that at least some aspects of co-localization of PRRs in the PM is mediated by *S*-acylation mediated by specific PAT proteins. As illustrated for FLS2, P2K1, and CERK1, examples of three, distinct families of RLKs, *S*-acylation is required for co-localization into PM nanodomains, which appears to coincide with an inactive, resting state for these receptors. Ligand activation of the receptors leads to a reduction in *S*-acylation, rapid autophosphorylation and subsequent turnover. In this way, *S*-acylation acts as a negative regulator of innate immunity, insuring that spurious activation does not occur, which might be detrimental to growth. Consistent with this model (Figure 6D), treatment with inhibitors of *S*-acylation or studies using mutants defective in PAT activity (*Atpat5*/*Atpat9*) or receptors mutated in their specific *S*-acylation sites all led to a marked elevation of the innate immunity response whether measured using cellular assays (e.g., cytoplasmic calcium levels) or pathogen sensitivity. A similar dynamic of receptor post-translational modification is also revealed during activation of the epidermal growth factor receptor (EGFR) in human tumorigenesis and cancer progression (Runkle et al., 2016). Hence, at least for these specific examples, the ‘yin-yang’ between *S*-acylation and phosphorylation/degradation is common across plants and animals.

In summary (Figure 6D), under steady-state conditions, plant PRRs are *S*-acylated and distributed into PM nanodomains in an inactive state. In response to ligand binding, the receptors are deacylated, altering their nanodomain localization, which occurs concomitantly with their autophosphorylation. This activation ultimately leads to receptor endocytosis and ubiquitin mediated degradation. PRR activation triggers downstream innate immune signaling, including rapid elevation of calcium influx, ROS, MAPKs activation and other events leading to a robust innate immunity response. In the case where the PRR is not degraded, *S*-acylation is restored as an alternative means to dampen the immune response. These results are important since they clearly demonstrate how *S*-acylation mediates plasma membrane machinery and signaling during plant innate immunity, drawing clear parallels with well-studied signaling systems in animals. Missing steps include identification of the specific thioesterases that mediate the rapid deacylation, as well as detailed molecular information regarding other components of both the inactive, *S*-acylated nanoclusters and those that likely form upon receptor activation. Hence, much remains to be discovered about how plants, as well as other organisms, are able to survive in the face of the variety and increasing severity of environmental change.

## ACKNOWLEDGMENTS

Research reported in this publication was supported by the National Institute of General Medical Sciences of the National Institutes of Health (grant no. R01GM121445 to G.S.). The content is solely the responsibility of the authors and does not necessarily represent the official views of the National Institutes of Health. This work was also supported by the Next-Generation BioGreen 21 Program Systems and Synthetic Agrobiotech Center, Rural Development Administration, Republic of Korea (grant no. PJ013254 to G.S.). Additional funding was provided within the framework of the 3^rd^ call of the ERA-NET for Coordinating Action in Plant Sciences through NSF grant 1826803 (to G.S.).

## AUTHOR CONTRIBUTIONS

D.C. designed and performed the experiments and wrote the manuscript. N.A. and J.J.T performed and supervised the kinase client screen for kinase targets. G.S. supervised the study and edited the manuscript. All authors discussed the results and commented on the manuscript.

## Methods

### CONTACT FOR REAGENT AND RESOURCE SHARING

Further information and requests for resources and reagents should be directed to and will be fulfilled by the Lead Contact, Gary Stacey.

### EXPERIMENTAL MODEL AND SUBJECT DETAILS

#### Plant Materials and Growth Conditions

All *Arabidopsis thaliana* plants used in this study are derived from the Columbia (Col-0) ecotype and express an T-DNA carrying aequorin (Knight and Knight, 1995), including *p2k1-3* (Salk-042209), *fls2* (SALK-026801), *cerk1* (GABI-KAT 096F09), *Atpat5* (GABI-322D08) and *Atpat9* (SALK-003020C). Additional transgenic lines in this study are described below. Plants were grown in soil or 1/2 MS medium containing 1% sucrose at 21-23 °C, 60-70% humidity in a growth chamber under long day (16 h light/8 h dark) conditions.

### METHOD DETAILS

#### Constructs and Transgenic Plants

Full-length CDS of P2K1 (At5g60300), FLS2 (AT5G46330), CERK1 (AT3G21630), PAT5 (At3g48760), PAT9 (At5g50020), RBOHC (AT5G51060), CD2b (At5g09390) and FLOT1 (AT5G25250), as well as their kinase domain or C-terminal domain, were amplified from wild-type plants using gene-specific primers (Table S1). The PCR products were cloned into pDONR-Zeo or pGEM-T Easy vectors. The different mutant forms were generated by PCR-mediated site directed mutagenesis.

In order to generate constructs for the LCI assays in tobacco leaves, the full-length DNA from the pDONR-Zeo vector were cloned into pCAMBIA1300-Nluc and pCAMBIA1300-Cluc using LR cloning. For BiFC assays, we used pAM-PAT-35SS::YFP:GW, pAM-PAT-35SS::YFPc:GW, and pAM-PAT-35SS::YFPn:GW as the destination vectors to form fusions with split YFP at the C-termini of proteins in *Arabidopsis* protoplasts using LR cloning. The pUC-GFP-GW and pUC-RFP-GW vectors fusing different mutated forms of P2K1, FLS2 and CERK1 were generated using LR cloning. These constructs were used for PM nanodomain localization in *Arabidopsis* protoplasts. The gateway plasmids pUC-GFP-GW and pUC-RFP-GW were generated by amplification of DNA fragments containing the 35S promoter and Nos terminator with SpeI/KpnI sites from pGWB5 and pGWB654 into prelinearized pUC19 digested by XbaI/KpnI, respectively.

To express specific proteins in *Arabidopsis* protoplasts, different mutated forms of P2K1, FLS2, CERK1, PAT5 and PAT9 from the source pDONR-Zeo source vectors were cloned into pUC-GW 14 and pUC-GW17 vectors using LR cloning.

In order to generate *NP::ATPAT5/Atpat5* and *NP::ATPAT9/Atpat9* stable complemented transgenic plants, the coding sequences driven by their native promoters were PCR-amplified from wild-type genomic DNA and cloned into pGWB14 using LR cloning.

In order to generate constructs used for expressing different mutant forms of P2K1 and CERK1, the native promoters of P2K1 and CERK1 about 1.5 kb before the start codon were amplified from genomic DNA by PCR, and used to replace the 35S promoter sequences of pFGC14, which was then fused with clones containing different *S*-acylation site forms of P2K1 and CERK1 CDS using LR cloning.

For constructs expressing recombinant proteins in *E. coli*, the DNA fragments of P2K1-KD, FLS2-KD and CERK1-KD cut with EcoRI and XhoI, EcoRI and XhoI, BamHI and XhoI were inserted into pGEX-5X-1, respectively. To gain His tagged constructs, DNA fragments of PAT5-CRD, PAT5-C, PAT9-CRD and PAT9-C cut with BamHI and XhoI were cloned into pET28a, while CD2b using SacI and XhoI, respectively.

#### Kinase client assay (KiC assay)

The KiC assay, instrument and detailed search parameters were used as before (Ahsan et al., 2013; Chen et al., 2017a). Two sets of empty vectors (GST and MBP) and two kinase-dead proteins, GST-P2K1-1 (D572N) and GST-P2K1-2 (D525N), were used as negative controls.

#### Calcium influx assay

Briefly, 4 or 5 days old seedlings were individually incubated with 50 μl of reconstitution buffer containing 10 μM coelenterazine (Nanolight Technology, Pinetop, AZ), 2mM MES buffer (pH 5.7), and 10 mM CaCl_2_ in the wells of a 96-well plate in the dark at room temperature overnight. The next morning, 50 μl of treatment solution (concentration was double strength to give a set final concentration of 25 mM MES and 100 μM ATP, 1 μM flg22, or 50 μg/ml chitin) was added to each well, and the luminescence was monitored using a CCD camera (Photek 216; Photek, Ltd.).

#### Oxidative (ROS) burst assay

Generally, leaf discs were taken from 4 or 5-week-old plants and incubated with 50 μl ddH_2_O in the wells of a 96-well-plate in the dark overnight. The next day, 50 μl 2 x chemiluminescent luminol buffer was added to each well to be a final concentration of 25 μM luminol, 20 μg/ml horseradish peroxidase and 200 μM ATPγS, 1 μM flg22, or 50 μg/ml chitin. Luminescence was immediately monitored using a CCD camera (Photek 216; Photek, Ltd.).

#### MAPK phosphorylation assay

Leaf discs from 4- or 5-week-old plants were incubated in the ddH_2_O overnight at 23°C, and then treated with 200 μM ATPγS, 1 μM flg22, or 100 μg/ml chitin for 0, 15, 30, or 60 min. Total protein was extracted with extraction buffer containing 50 mM Tris (PH 7.5), 150 mM NaCl, 0.5% Triton-X 100, and 1 ^x^ protease inhibitor for 30 min on ice. The extracted total proteins were separated by 10% (w/v) SDS-PAGE gel and detected by immunoblotting with anti-phospho-p44/p42 MAPK antibody (dilution, 1:1000).

#### Bacterial growth assays

Bacterial growth was performed using flood inoculation of seedlings. Generally, 50 ml of *P. syringae pv. tomato* DC3000 (5 x 10^6^ colony-forming units (CFU) ml^−1^) bacterial suspension containing 10 mM MES pH 5.7, 10 mM MgCl_2_, 0.025% Silwet L-77 was dispensed into plates containing about 14-day-old seedlings for 2-3 min. After 2 days post inoculation, the roots of seedlings were removed and the seedlings were washed with ddH_2_O for 5 min. The seedlings without roots were ground in 10 mM MgCl_2_, diluted serially, and plated on LB agar with 25 mM rifampicin. Colonies (CFU) were counted after incubation at 28°C for 2 days.

#### Split-luciferase complementation imaging (LCI) assay

The *Agrobacterium tumefaciens* (GV3101) cells containing the indicated constructs were infiltrated into 4-week-old leaves of *N. benthamiana* and then incubated at room temperature for 48 hours before LUC activity measurement. 1 mM D-luciferin was sprayed onto the leaves and then kept in the dark for 5-10 min to allow the chlorophyll luminescence to decay, the luminescence was monitored using a CCD camera (Photek 216; Photek, Ltd.).

#### *Arabidopsis* protoplast isolation and transformation

The isolation and transfection of *Arabidopsis* protoplasts were performed as previously described (Chen et al., 2017a) and used for the various assays as indicated.

#### Bimolecular fluorescence complementation (BiFC) assay

N- and C-terminal YFP protein fusions plasmids were co-transformed into *Arabidopsis* protoplasts as described above and then incubated at 23°C in a growth chamber overnight at dark. The YFP fluorescence was observed using a Leica DM 5500B Compound Microscope with Leica DFC290 Color Digital Camera. FM4-64 was used as the plasma membrane stain.

#### *In vitro* pull-down

Recombinant proteins GST-P2K1-KD, GST-P2K1-KD-1, GST-FLS2-CD, GST-CERK1-C, His-PAT5-CRD, His-PAT5-C, His-PAT9-CRD, His-PAT9-C, or His-CD2b were expressed in *E. coli* and affinity purified using Glutathione Resin (GenScript) and TALON^®^ Metal Affinity Resin (Clontech), respectively. For pull-down, 5 μg GST and His recombinant proteins were incubated with 25 μl Glutathione Resin beads in the pull down buffer containing 25mM Tris-HCl pH 7.5, 100 mM NaCl, and 1 mM DTT for 2 h at 4 °C. The beads were washed more than seven times with the washing buffer 25mM Tris-HCl pH 7.5, 100mM NaCl, and 0.1% Triton-X 100. The bund proteins were eluted with 25 μl elution buffer containing 50 mM Tris-HCl pH 7.5-8, 15-20 mM GSH for about 15-30 min. The proteins were separated using SDS-PAGE gels and detected by immunoblotting using anti-His (dilution, 1:1000) and anti-GST-Hrp (dilution, 1:1000).

#### *In vitro* phosphorylation assays

For the *in vitro* kinase assay, 2 μg of purified GST-P2K1-KD, GST-P2K1-KD-1, GST-FLS2-CD or GST-CERK1-KD kinases were incubated with 1 μg His-PAT5-CRD, His-PAT5-C, His-PAT9-CRD, His-PAT9-C, or His-CD2b as substrate in a 20 μl reaction buffer containing 50 mM Tris-HCl pH 7.5, 50 mM KCl, 10 mM MgCl_2_, 10 mM ATP, and 0.25 μl radioactive [γ-^32^P] ATP for 30 min at 30°C. The reaction was stopped by 5 μl of 5 x SDS loading buffer. The proteins were separated by SDS-PAGE (10%), followed by autoradiography for 3 h. The proteins within the gel were visualized by staining with Coomassie blue. Myelin basic protein (MBP) was used as a positive control.

#### Co-immunoprecipitation assay

Total proteins were extracted from protoplasts or plant tissues with an extraction buffer containing 50 mM Tris (PH 7.5), 150 mM NaCl, 0.5% Triton-X 100, and 1 x protease inhibitor for 1 h on ice. The samples were centrifuged at 20,000 x g for 15 min at 4°C, the supernatant was decanted and 1 μg anti-Myc or 30 μl anti-HA agarose was added to the supernatant and incubated for 4 h or overnight with end-over-end shaking at 4°C. 25 μl protein A resin was added for 2 h, spun down and washed seven times with extraction buffer. After washing, 25 μl 1 x SDS-PAGE loading buffer was added and heated at 100°C for 10 min. The proteins were separated by SDS-PAGE and detected by immunoblotting with anti-HA-HRP (dilution, 1:3000), anti-Myc-HRP (dilution 1:3000) anti-FLS2 (dilution, 1:5000), or anti-CERK1 (dilution, 1:4000).

#### *S*-Acylation Assay

*Arabidopsis* protoplasts transfected with HA-tagged proteins were homogenized in lysis buffer containing 50 mM Tris (PH 7.5), 150 mM NaCl, 0.5% Triton-X 100, and 1 x protease inhibitor for 1 h on ice. After centrifuged at 20,000 x g for 15 min at 4°C, 50 mM N-ethylmaleimide was added to the supernatant for blocking free sulfhydryl groups, and proteins were then immunoprecipitated using Anti-HA-Agarose beads with end-over-end shaking at 4°C overnight. The next day, the beads were washed three times with lysis buffer and eluted in 100 μl 0.1 μg/μl HA peptide for 15 min. The eluted proteins were divided into two equal portions: one treated with 1 M hydroxylamine and the other with 1 M Tris.HCl (pH 7.4) (as a control) in the presence of activated thiol-Sepharose 4B. Two hours later, the sepharose beads were washed three times with lysis buffer without protease inhibitor at room temperature, and then resuspended in 1 x protein loading buffer and heated at 100°C for 10 min. Western blots were performed using anti-HA-HRP (dilution, 1:3000).

#### Confocal laser scanning microscopy and nanodomain detection

The *Arabidopsis* protoplasts after transformation were imaged on a Leica SP8 spectral confocal microscope using a 63 x water immersion lenses or 100 x oil immersion objectives. The GFP was excited with solid-state laser diode at 488 nm and the emission detected at 505-550 nm, and the RFP was excited with solid-state laser diode at 558 nm and the emission detected at 570-650 nm. The fluorescence images obtained with a Leica CCD camera.

For nanodomain detection, the microscopic images of GFP and RFP from the same protoplast and taken at the same time were exported to Huygens Professional software for deconvolution. After verified by Huygens Professional, the images were followed the deconvolution wizard process and were exported back to Leica LAS X software.

For quantitative co-localization analysis, a region of interest (ROI) was selected and quantified using the colocalization tool within the Leica LAS X software. All ROI were of the same size, contrast and brightness.

### QUANTIFICATION AND STATISTICAL ANALYSIS

Error bars in the figures are standard deviation (SD) or the standard error of the mean (SEM = SD/(square root of sample size)), and number of replicates is reported in the figure legends. Statistical comparison among different samples was carried out by one-way ANOVA. Samples with statistically significant differences (*p < 0.05 or **p < 0.01 as indicated in the figure legends) are marked with different letters (a, b, c etc.).

### DATA AND SOFTWARE AVAILABILITY

The GSP-Lipid 1.0 software is online: http://lipid.biocuckoo.org/. Huygens Professional is: https://svi.nl/Huygens-Professional.

