## Supplemental Figures for "*S*-Acylation of plant immune receptors mediates immune signaling in plasma membrane nanodomains"

### SUPPLEMENTAL INFORMATION

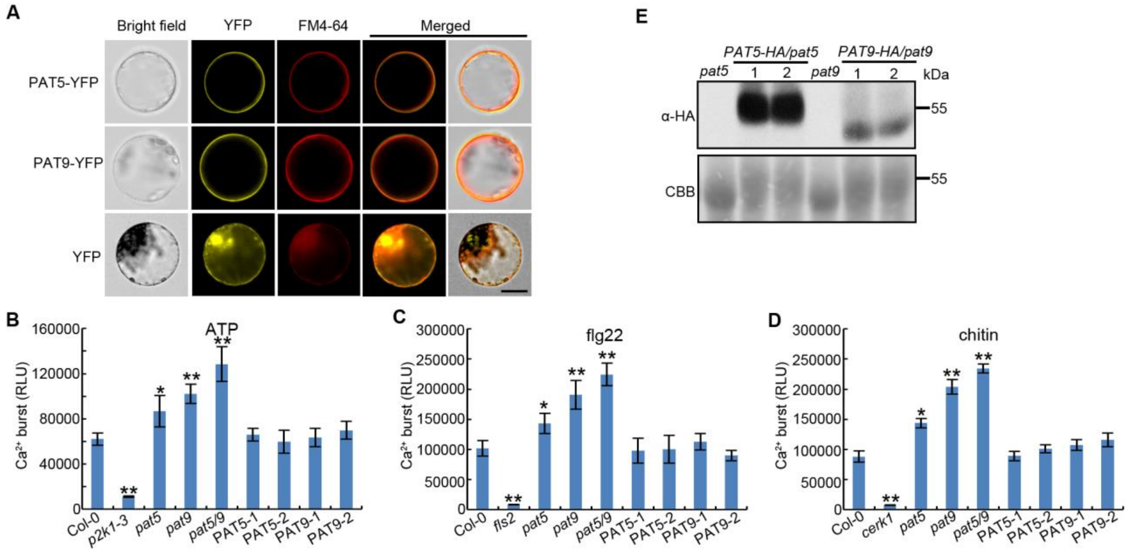


**Figure S1. Subcellular localization of PAT5 and PAT9, Related to Figure 1**

(A) PAT5-YFP and PAT9-YFP were transiently expressed in the *Arabidopsis* protoplasts, and merged well with the plasma membrane marker FM-64. Free YFP was used a control. Bar = 20 μm.

(B-D) Ligand-induced calcium influx. 5-day-old seedlings were treated with 100 μM ATP, 1 μM flg22 or 50 μg/ml chitin. RLU, relative luminescence units.

(E) Relative expression protein levels of *NP::ATPAT5-HA/Atpat5* and *NP::ATPAT9-HA/Atpat9* complemented transgenic lines using their own native promoters. CBB was used as a loading control.

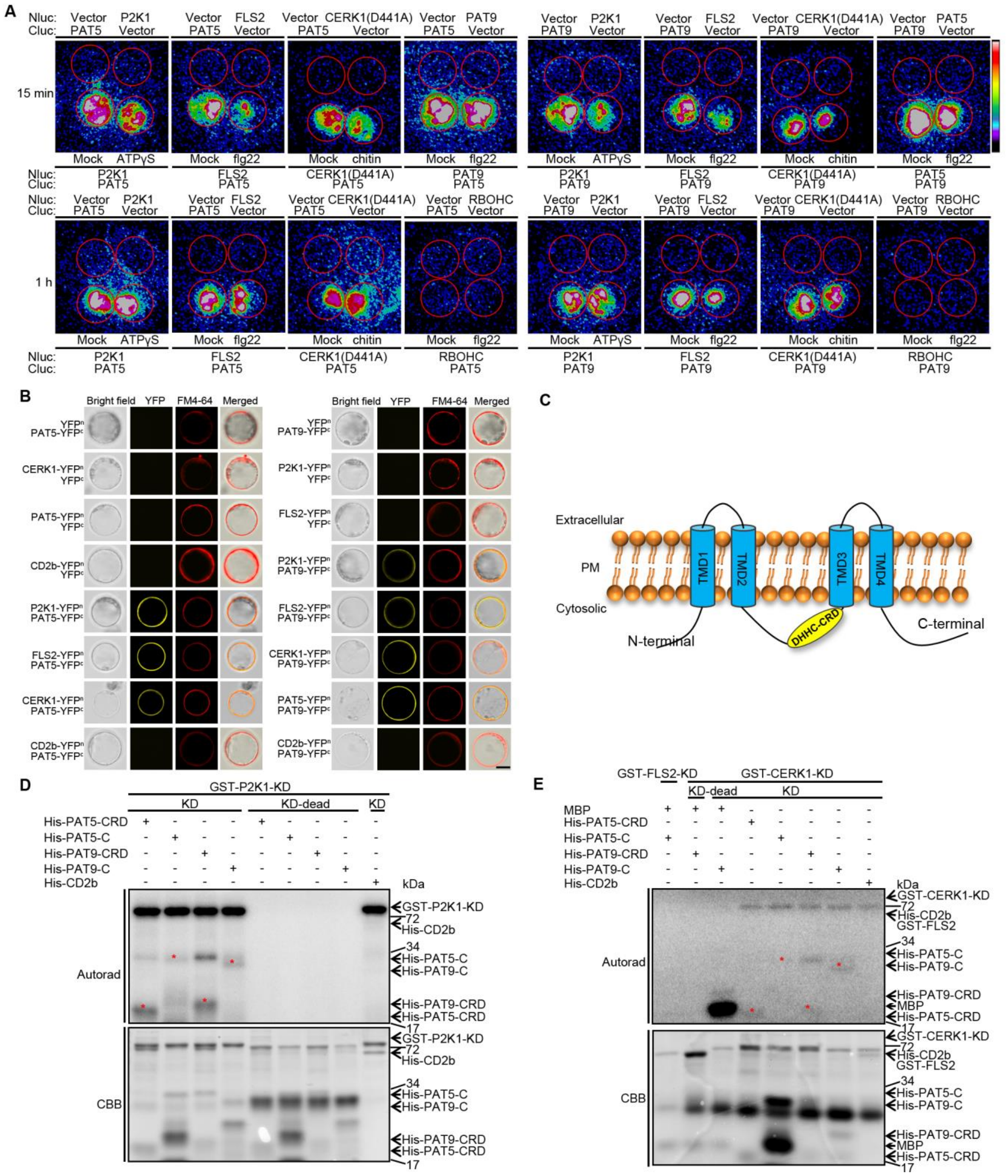

**Figure S2. PAT5 and PAT9 interact with and are phosphorylated by receptors, Related to Figure 2**

(A) PAT5 and PAT9 interact with P2K1, FLS2 and CERK1 in *N. benthamiana* in an elicitor-dependent manner. The indicated constructs were transiently expressed in *N. benthamia* leaves with, where indicated in red circles, the addition of 500 μM ATPγS, 2 μM flg22 or 100 μg/ml chitin. Nluc, N-terminal fragment of firefly luciferase; Cluc, C-terminal fragment of firefly luciferase; Vector, empty vector. The color bar represents the color code for fluorescence intensities.

(B) Interactions of PATs and receptors at *Arabidopsis* protoplast plasma membrane. FM4-64 was used to stain the plasma membrane. Bar = 20 μm.

(C) Protein topology of PATs on plasma membrane. PM, plasma membrane. TMD, transmembrane domains. DHHC, Asp-His-His-Cys. CRD, a stretch of DHHC within a Cys-rich domain.

(D and E) Receptors directly phosphorylate PAT5 and PAT9. Purified P2K1, FLS2 and CERK1 kinase domain recombinant proteins were incubated with PAT5/PAT9-CRD and -C domains in an *in vitro* kinase assay. Autophosphorylation and trans-phosphorylation were measured by incorporation of γ-[32P]-ATP. MBP and GST-CD2b were used as positive and negative controls, respectively. The protein loading was measured by CBB staining. Red stars represent the trans-phosphorylated proteins.

**A**

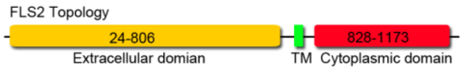


| ID | Position | Peptide | Score | Cutoff | Cluster |
|---|---|---|---|---|---|
| FLS2 | 61 | IIGSLRHCNWTGITC | 0.195 | 0.0 | S-Palmitoylation: Cluster C |
| FLS2 | 68 | CNWTGITCDSTGHVV | 0.293 | 0.0 | S-Palmitoylation: Cluster B |
| FLS2 | 165 | GDVPEEICKTSSLVL | 0.835 | 0.0 | S-Palmitoylation: Cluster B |
| FLS2 | 187 | LTGKIPECLGDLVHL | 0.499 | 0.0 | S-Palmitoylation: Cluster A |
| FLS2 | 263 | IPAEIGNCSSLVQLE | 0.018 | 0.0 | S-Palmitoylation: Cluster A |
| FLS2 | 407 | IPSSISNCTGLKLLD | 0.0 | 0.0 | S-Palmitoylation: Cluster C |
| FLS2 | 454 | IPDDIFNCSNLETLS | 0.244 | 0.0 | S-Palmitoylation: Cluster A |
| FLS2 | 672 | IPRSLQACKNVFTLD | 0.447 | 0.0 | S-Palmitoylation: Cluster C |
| FLS2 | 783 | LMGNTDLCGSKKPLK | 0.783 | 0.0 | S-Palmitoylation: Cluster C |
| FLS2 | 792 | SKKPLKPCTIKQKSS | 0.0 | 0.0 | S-Palmitoylation: Cluster B |
| FLS2 | 830 | LLVLILTCCKKKEKK | 9.137 | 0.903 | S-Palmitoylation: Cluster C |
| FLS2 | 831 | LVLILTCCKKKEKKI | 8.015 | 0.903 | S-Palmitoylation: Cluster C |
| FLS2 | 975 | LLEKIDLCVHIASGI | 2.121 | 1.991 | S-Palmitoylation: Cluster B |
| FLS2 | 996 | YGFPIVHCDLKPANI | 0.257 | 0.0 | S-Palmitoylation: Cluster C |
| FLS2 | 1132 | IEDFLKLCLFCTSSR | 0.795 | 0.0 | S-Palmitoylation: Cluster C |
| FLS2 | 1135 | FLKLCLFCTSSRPED | 0.329 | 0.0 | S-Palmitoylation: Cluster C |

**B**

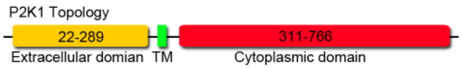


| ID | Position | Peptide | Score | Cutoff | Cluster |
|---|---|---|---|---|---|
| P2K1 | 87 | SFSTHFVCALVPKPG | 0.447 | 0.0 | S-Palmitoylation: Cluster B |
| P2K1 | 296 | LIILLPVCLAILVLA | 0.502 | 0.0 | S-Palmitoylation: Cluster A |
| P2K1 | 394 | AEVVSMRCLKHRNLV | 2.301 | 1.991 | S-Palmitoylation: Cluster B |
| P2K1 | 407 | LVPLFGYCRRKRELL | 1.966 | 0.903 | S-Palmitoylation: Cluster C |
| P2K1 | 538 | VFMLEVTCGRRPVEP | 0.461 | 0.0 | S-Palmitoylation: Cluster C |
| P2K1 | 559 | RHMIKWVCECWKKDS | 1.598 | 0.0 | S-Palmitoylation: Cluster B |
| P2K1 | 561 | MIKWVCECWKKDSLL | 0.638 | 0.0 | S-Palmitoylation: Cluster C |
| P2K1 | 594 | VMKLGLLCSNIVPES | 0.355 | 0.0 | S-Palmitoylation: Cluster A |

**C**

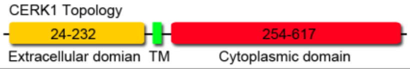


| ID | Position | Peptide | Score | Cutoff | Cluster |
|---|---|---|---|---|---|
| CERK1 | 25 | FFAVESKCRTSCPLA | 3.048 | 1.991 | S-Palmitoylation: Cluster B |
| CERK1 | 29 | ESKCRTSCPLALASY | 1.817 | 0.893 | S-Palmitoylation: Cluster A |
| CERK1 | 91 | RVLVPFPCECQPGDF | 0.701 | 0.0 | S-Palmitoylation: Cluster C |
| CERK1 | 93 | LVPFPCECQPGDFLG | 0.684 | 0.0 | S-Palmitoylation: Cluster C |
| CERK1 | 153 | TLNVLVNCSCGDESV | 0.038 | 0.0 | S-Palmitoylation: Cluster C |
| CERK1 | 155 | NVLVNCSCGDESVSK | 0.136 | 0.0 | S-Palmitoylation: Cluster C |
| CERK1 | 381 | LVRLIGYCVEGSLFL | 0.328 | 0.0 | S-Palmitoylation: Cluster C |
| CERK1 | 572 | MAELGKACTQENAQL | 0.943 | 0.0 | S-Palmitoylation: Cluster B |

**Figure S3. FLS2, P2K1 and CERK1 protein domains and *S*-acylation sites prediction, Related to Figure 3**

(A-C) Schematic representation of FLS2, P2K1 and CERK1 protein structures contain an extracellular domain, a transmembrane domain (TM) and a cytoplasmic domain. The numbers in the topology and position indicate amino acids. The receptor protein sequences were analyzed by GPS-Lipid 1.0 software, the *S*-acylation residues were shown in red font.

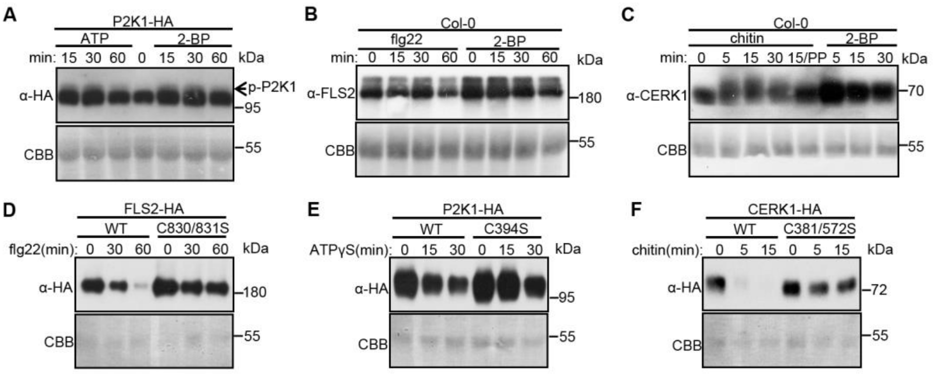


**Figure S4. PAT5 and PAT9 regulate P2K1, FLS2 and CERK1 phosphorylation and degradation, Related to Figure 4**

(A) P2K1-HA phosphorylation was analyzed by immunoblot in wild type plant upon addition of 200 μM ATPγS or 50 μM 2-BP (*S*-acyltransferase inhibitor). p-P2K1, phosphorylation of P2K1.

(B) 2-BP reduced endogenous FLS2 degradation. Leaf discs of Col-0 treated with 20 μM flg22 and 50 μM 2-BP were used for FLS2 protein detection.

(C) 2-BP enhances endogenous CERK1 phosphorylation and reduces CERK1 degradation. Leaves of Col-0 infiltrated with 100 μg/ml chitin and 50 μM 2-BP were used for CERK1 protein detection. PP, lambda protein phosphatase.

(D-F) Analysis of the rate of turnover of P2K1, FLS2 and CERK1 proteins modified (C→S) at the site of *S*-acylation. The indicated constructs were transformed into *Arabidopsis* wild-type Col-0 protoplasts treated with 200 μM ATPγS, 10 μM flg22 or 100 μg/ml chitin.

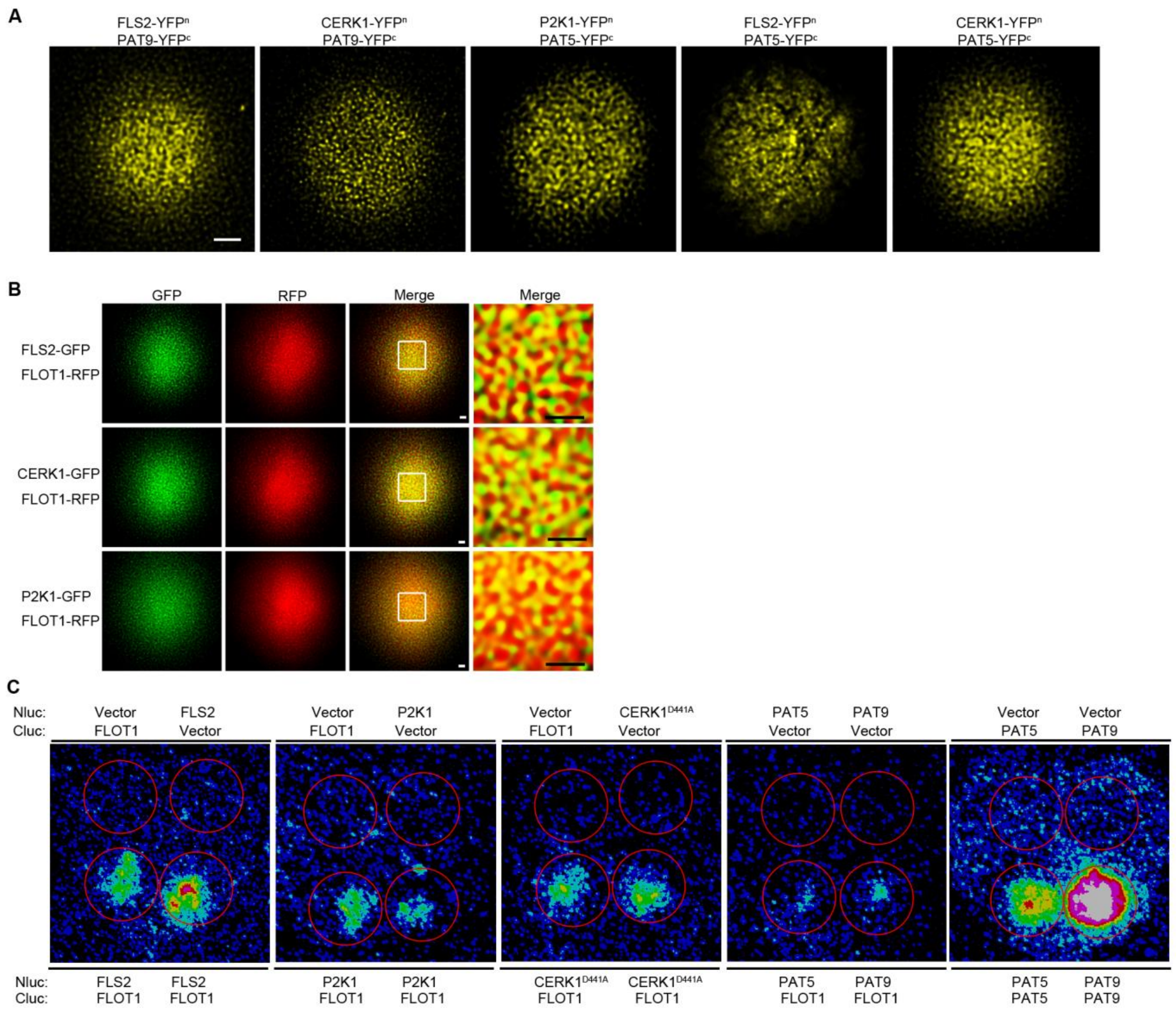


**Figure S5. PAT5, PAT9 and FLOT1 interact with FLS2, CERK1 and P2K1 in PM nanodomains, Related to Figure 5**

(A) PAT9 and PAT5 interact with receptors in PM nanodomains. The indicated constructs were transiently co-expressed in *Arabidopsis* protoplasts, the dispersed punctate fluorescence signaling was observed and analyzed using confocal microscopy. Bar = 5 μM.

(B) FLS2, CERK1 and P2K1 co-localize with the nanodomain marker flotilin (FLOT1). Confocal micrographs of FLS2-, CERK1- and P2K1-GFP plasma membrane localization after transient co-expression with FLOT1-RFP in *Arabidopsis* protoplasts. The dashed squares represent the areas magnified within the far right image. Bar = 1 μM.

(C) FLOT1 interacts with FLS2, CERK1, P2K1, PAT5 and PAT9 by the firefly LCI assays in tobacco leaves in the dark. PAT5 and PAT9 form homodimers and show stronger LCI signaling.

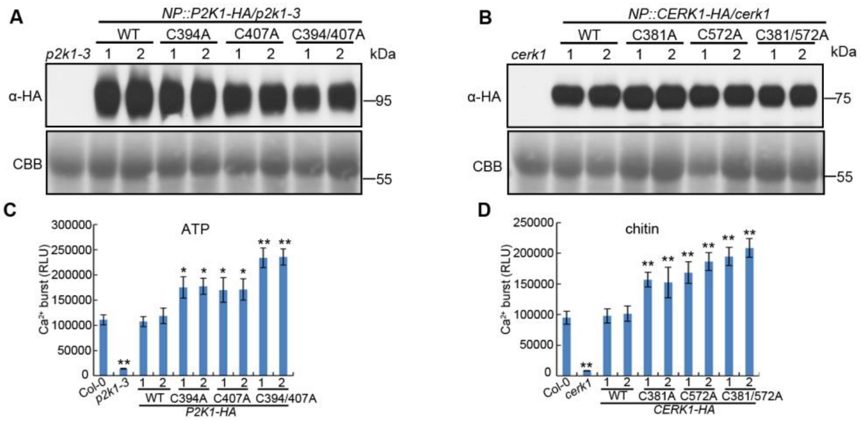


**Figure S6. *S*-Acylation of FLS2, CERK1 and P2K1 negatively mediate PTI, Related to Figure 6**

(A and B) Total P2K1-HA and CERK1-HA proteins were detected by anti-HA immunoblot for the stable transgenic plants. CBB, Coomassie brilliant blue staining.

(C and D) ATP and chitin-induced calcium influx were significantly increased in plants expressing CERK1 or P2K1 proteins lacking critical cysteine residues for *S*-acylation. RLU, relative luminescence units; Error bars indicate ± SEM; n = 12 (biological replicates, one-sided ANOVA). $^{*}P < 0.05$, $^{**}P < 0.01$, Student’s t test.
